# The prefoldin-like protein AtURI exhibits characteristics of instrinsically disordered proteins

**DOI:** 10.1101/2023.12.13.571423

**Authors:** Yaiza Gómez-Mínguez, Alberto Palacios-Abella, Cecilia Costigliolo-Rojas, Mariana Barber, Laura Hernández-Villa, Cristina Úrbez, David Alabadí

**Affiliations:** Instituto de Biología Molecular y Celular de Plantas (CSIC-UPV), Valencia (Spain); Fundación Instituto Leloir, Buenos Aires (Argentina)

**Author notes:** For correspondence. Mailing address: IBMCP (CSIC-UPV), Edificio E8, Campus UPV, Ingeniero Fausto Elio s/n, 46022-Valencia (Spain).

**Keywords:** protein disorder, interactome, protein stability

## Abstract

The prefoldin-like protein UNCONVENTIONAL PREFOLDIN RPB5 INTERACTOR (URI) participates in diverse cellular functions, including protein homeostasis, transcription, translation, and signal transduction. Thus, URI is a highly versatile protein, although the molecular basis of this versatility remains unknown. In this work, we show that *Arabidopsis thaliana* (*Arabidopsis*) URI (AtURI) possesses a large intrinsically disordered region (IDR) spanning most of the C-terminal part of the protein, a feature conserved in yeast and human orthologs. Our findings reveal two key characteristics of disordered proteins in AtURI: promiscuity in interacting with partners and protein instability. We propose that these two features contribute to providing AtURI with functional versatility.

## INTRODUCTION

Animal and yeast models show that prefoldin-like proteins are regulators of cell survival and proliferation in response to environmental and nutrient stress [1]. Prefoldin-like proteins were identified in several independent studies in animals and yeast. They are UNCONVENTIONAL PREFOLDIN RPB5 INTERACTOR (URI; Bud27 in yeast) [2], UBIQUITOUSLY EXPRESSED TRANSCRIPT (UXT) [3], p53 AND DNA DAMAGE-REGULATED GENE1 (PDRG1) [4] and ASNSD1 UPSTREAM READING FRAME (ASDURF) [5]. More recently, the *Arabidopsis* ortholog of URI, AtURI, has been identified as a novel regulator of auxin signaling [6]. Sequence analysis showed that they are related to the canonical prefoldins, which form the prefoldin (PFD) complex, a cochaperone previously identified in humans and yeast [7,8]. Proteomic approaches in human cell lines have shown that URI, UXT, PDRG1 and ASDURF, together with the canonical prefoldins PFD2 and PFD6 form the prefoldin-like (PFDL) complex, which is probably involved in protein homeostasis [2,5,9]. This complex does not exist in yeast, since URI/Bud27 is the only prefoldin-like protein found in this microorganism.

URI is unique among prefoldin and prefoldin-like proteins in that it is a larger protein. In addition to its involvement in several cellular functions related to its role as a cochaperone, either as part of the PFDL complex or as an independent subunit, its role as a signaling protein has also been described [1]. In yeast, URI/Bud27 is involved in transcription, indirectly through the assembly of the three nuclear RNA polymerases in the cytoplasm [10] and directly through the assembly of RNA polymerase II transcription complexes at the chromatin [11,12]. URI plays a role as a signaling protein in the context of nutrient signaling in humans. Specifically, URI phosphorylation prevents its interaction with the phosphatase PP1γ, an event that triggers different signaling cascades depending on the energy status of the cell [13,14]. Recently, the first study on the role of URI in plants was reported. In *Arabidopsis*, AtURI is involved in the intracellular transport of the auxin efflux carrier PIN-FORMED2, which is necessary to establish the asymmetry of auxin accumulation in the root to respond to gravitropic stimuli [6]. Authors suggested that the involvement of AtURI in auxin transport could be mediated by the role of URI in cytoskeleton assembly. The prefoldin-like URI is therefore a versatile protein participating in various cellular processes that involve protein homeostasis and signal transduction. However, the molecular basis for this versatility is not known.

Many functional proteins are completely unstructured or contain unstructured regions, i.e., regions that cannot fold into a well-defined structure [15]. These are referred to as instrinsically disordered proteins and instrinsically disordered regions (IDR). Structural disorder is important for the function of many regulatory proteins acting in signaling cascades and in processes involving the assembly of macromolecular complexes [16]. For example, the flexibility of IDRs makes it easier for proteins to interact with a variety of partners or to undergo post-translational modifications. Analyses of protein-protein interaction networks in various organisms showed that proteins that establish more interactions than average have a tendency to be disordered [17,18]. In addition, disordered proteins and proteins with IDRs tend to be unstable, so their amount can be tightly controlled, which is an advantageous property for a regulatory protein [19], preventing potential adverse effects of high amounts of these proteins [20]. In this work, we show that AtURI possesses properties of intrinsically disordered proteins. These properties likely contribute to the functional versatility of AtURI, making it the type of regulatory protein in signaling pathways and protein homeostasis.

## MATERIALS AND METHODS

### Protein affinity purification in *Arabidopsis* cell suspensions

The coding sequences (CDSs) of *AtURI1* and *UXT* in *pENTR223* were obtained from the Arabidopsis Biological Resource Center (OH, USA), and were transferred into the *pKNGS_rhino* vector [21] using Gateway. *Agrobacterium tumefaciens* C58 cells carrying *pKNGS_rhino-AtURI1*, *pKNGS_rhino-UXT* and the control construct *pKNGS_rhino* were used to transform PSB-D cell cultures [21]. The preparation of protein extracts and the one-step affinity purification (AP) of GS-AtUR1, GS-UXT and GS alone were carried out as previously described [22]. Proteins in the purified samples were identified by LC-MS/MS at the Unidad de Proteómica (Universidad de Córdoba, Spain).

### In silico analyses

The sequences used for phylogenetic analysis of UXT and ASDURF were acquired through TBLASTN analysis for each species. The data sources utilized were the Phytozome database (https://phytozome-next.jgi.doe.gov/) and NCBI (https://www.ncbi.nlm.nih.gov/). The amino acid sequences of the proteins were used to construct a maximum likelihood (PhyML) phylogenetic tree using default parameters with NGPhylogeny.fr [23]. This tree included UXT and ASDURF sequences from *Oryza sativa*, *Arabidopsis thaliana*, *Medicago truncatula*, *Vitis vinifera*, *Aquilegia coerulea*, *Musa acuminata*, *Mus musculus*, and *Homo sapiens*. PFD1 and PFD5 from *Arabidopsis* were used as outliers. To ensure a phylogenetic tree with stronger support, sequences with insertions or missing bases were eliminated from the final alignment.

The 3D structure of each *Arabidopsis* prefoldin-like protein was determined by homology with the models of *Arabidopsis* canonical prefoldins [24] using Modeller (release 10.1) [25]. Similarly, the 3D structure of each human prefoldin-like protein was determined by homology with the human canonical prefoldins (PDB code 6NR8) [26]. The 3D structure of the *Arabidopsis* and human PFDL complex was assembled using the human PFD complex (PDB code 6NR8) as a template and visualized using PyMOL 2.4 software.

The protein disorder was predicted with the online tools DEPICTER (http://biomine.cs.vcu.edu/servers/DEPICTER/) [27] and MobiDB (https://mobidb.bio.unipd.it/) [28], and with preFold (https://github.com/aretasg/preFold), a tool written in Python and inspired by FoldIndex [29].

### Yeast two-hybrid assays

The CDSs of *PFD2*, *PFD6*, and *PDRG1* in *pENTR223* were obtained from the Arabidopsis Biological Resource Center (OH, USA). *PFD2*, *PFD6*, *PDRG1*, and *UXT* CDSs were transferred into the *pGADT7-GW* destination vector to generate fusions with the Gal4 activation domain, while the *AtURI* CDS was transferred into *pGBKT7-GW* to generate a GAL4 DNA binding domain fusion by Gateway (Thermo Fisher Scientific). pGADT7- and pGBK7-derived expression vectors were introduced into Y187 and Y2HGold yeast strains, respectively. Protein interactions were tested in diploids by the nutritional requirement of histidine (His).

### Protein co-immunoprecipitation assays and Western blot analysis

HA-AtURI1 and YFP-UXT were prepared by transferring the CDSs into the *pEarleyGate201* and *pEarleyGate104* vectors [30], respectively, by Gateway and used to transfrom *Agrobacterium tumefaciens* C58. Leaves of one-month-old *Nicotiana benthamiana* plants grown under 16 hours light: 8 hours dark at 25°C were infiltrated with different mixtures of *Agrobacterium* cells carrying the expression vectors and the p19 silencing suppressor. Leaves were harvested three days after infiltration and frozen in liquid nitrogen. The preparation of proteins extracts and the co-immunoprecipitation analysis were performed as we previously described [31].

The CDSs of *AtURI^mut4A^* and *AtURI^mut5A^* were synthesized by Intregrated DNA Technologies (Belgium) and transferred into the *pEarleyGate104* vector by Gateway to create YFP fusions. The WT and the two mutant versions were transiently expressed in *N. benthamiana leaves* as explained above. Protein levels were analyzed by Western blot.

For Western blot analysis of protein levels from *Arabidopsis* or *N. benthamiana* samples, protein extracts were prepared as for co-immunoprecipitation. Protein samples were mixed with 1/4 volume of 4x Laemmli buffer and heated at 95°C for 5 minutes. Western analysis was carried out as above. In addition to anti-GFP and anti-HA-HRP antibodies, an anti-FLAG-HRP antibody (A8592, Sigma-Aldrich) and Peroxidase anti-Peroxidase Soluble Complex antibody (P1291, Sigma) were also used.

### Preparation of *Arabidopsis* transgenic lines

To prepare the *pAtURI:AtURI1-3xFLAG* construct, a genomic fragment starting 1312 nucleotides upstream of the *AtURI* ATG until the codon before the stop was amplified by PCR and a *pENTR207-gAtURI* was generated by Gateway. *gAtURI* was transferred into pEarleyGate302 containing a 3xFLAG tag [32] by Gateway. This construct and the *pEarleyGate104-AtURI* construct to overexpress *YFP-AtURI* were introduced into Col-0 WT *Arabidopsis* plants by *Agrobacterium*-mediated transformation.

### Production of recombinant MBP-AtURI in *E. coli*

The MBP-AtURI1 fusion was prepared by transferring the *AtURI1* CDS into the *pDEST-MBP-GW* vector [33] by Gateway and used to transform *E. coli* BL21 Rosetta cells. To induce protein expression, 0.5 mM IPTG was added into the culture at an OD_600_ of 0.53 and cells were incubated at 16°C for 16 hours. Cells were collected by centrifugation at 4500 rpm for 15 minutes. The cell pellet was resuspended in 5 mL of column buffer [20 mM Tris-HCl, 200 mM NaCl, 1mM EDTA, 1X protease inhibitor cocktail (cOmplete, EDTA-free; Roche), pH 7.4] and sonicated using a cell disruptor (Fisherbrand™ Model 120 Sonic Dismembrator). The sonicated sample was centrifuged at 14000 rpm for 20 min at 4°C. The supernatant was diluted with column buffer to a final volume of 35 mL and was filtered through a 0.45 μm membrane. One mL of amylose resine (E8021S, New England Biolabs) was poured into a 1.5 x 10 cm column. The column was washed with 5 column volumes (CVs) of column buffer. The diluted crude extract was loaded and the non-specifically bound proteins were washed out from the column with 10 CVs of column buffer. MBP-AtURI1 protein was eluted with 5 CV of column buffer supplemented with maltose 10 mM. Glycerol was added to the eluted fractions up to 20%. MBP-AtURI1 protein was stored at -80°C.

### Cell free protein degradation assays

Protein extracts from *N. benthamiana* leaves or *Arabidopsis* seedlings were prepared in extraction buffer (25 mM Tris-HCl pH 7.5, 10% glycerol, 1mM EDTA pH 8.0, 150 mM NaCl). The degradation assays were done in a volume of 250 μL containing 250 μg of total proteins. Protein extracts were supplemented with 0.5 mM cycloheximide (CHX; Merck), and the mixtures were then divided in two parts, one was supplemented with 150 μM MG-132 (Insolution™, Merck) or 100 μM PYR-41 (Sigma) and the other with 2% DMSO as a control. When MBP-AtURI was used, 10 ng of recombinant protein was added to the *Arabidopsis* protein extract. The mixtures were incubated at 30°C and 25 μL aliquots were taken at the indicated times. Incubations were stopped by adding 2x Laemli buffer to the aliquots and incubating 5 minutes at 95°C. Protein levels were analyzed by western blot and the band intensity quantified using ImageJ.

### RNA extraction and real-time qPCR

We used one-week-old WT and *YFP-AtURI Arabidopsis* seedlings grown under continuous white fluorescent light (60 μmol m^-2^s^-1^) at 22°C and inflorescences of 25-day-old plants of the same lines grown under 16 hours light: 8 hours dark at 22°C. Total RNA was isolated using the NucleoSpinTM RNA Plant Kit (Macherey-Nagel) following the manufacturer’s instructions. cDNA was prepared from 1 μg of total RNA with NZY-First Strand cDNA Synthesis Kit (NZYTech) according to the manufacturer’s instructions. The resulting cDNA was used for real-time quantitative PCR reaction. The primers used were: *AtURI* (CGTTATGGTTCCATTCGGTAAAA, TCTCCCAACAACACCAAACACT), *GFP* (ACCTACGGCAAGCTGACCC, CGGGCATGGCGGACTTGAAG), and *PP2AA3* (*At1g13320*; TAACGTGGCCAAAATGATGC, GTTCTCCACAACCGCTTGGT) as control gene.

## RESULTS AND DISCUSSION

### AtURI is part of the Prefoldin-like complex in *Arabidopsis*

In animals, URI forms the PFDL complex with the prefoldin-like proteins UXT, PDRG1, and ASDURF and the canonical prefoldins PFD2 and PFD6 [2,5,9]. In *Arabidopsis*, AtURI interacts with PFD2 and PFD6, as shown by yeast 2-hybrid (Y2H) and by co-immunoprecipitation assays [6]. Nevertheless, it remains unknown whether a PFDL complex exists in *Arabidopsis* and AtURI is one of its subunits. To investigate this possibility, we identified the AtURI interactors *in vivo* in *Arabidopsis* PSB-D cell suspensions [21]. We fused the GS tag to the N-terminal end of AtURI, which consists of a protein G-tag and the streptavidin-binding peptide [21]. Although the GS tag was designed for a two-step affinity purification, we performed a single immunoprecipitation to increase the chance of identifying weak or transient interactions, as previously described [22]. We prepared transgenic PSB-D cell suspensions expressing the unfused GS tag or GS-AtURI (Fig. S1) and extracts from two and three biological replicates, respectively, were subjected to an affinity purification (AP) step using mouse IgG-coated paramagnetic beads. Proteins in the eluates were identified by mass spectrometry (MS) analysis. We only considered proteins that were present in the three replicates, with minimum of two unique peptides in at least one of them, and that were not present in the GS control. Among the top interactors, we identified PFD2 and PFD6 in the eluates of GS-AtURI (Table 1; Table S1). We also identified the putative ortholog of the prefoldin-like protein PDRG1 (encoded by the *At3g15351* gene) and two proteins annotated as “Prefoldin chaperone subunit family protein”, encoded by *At1g26660* and *At1g49245* genes. None of these proteins were identified in the eluates from the control GS (Table 1; Table S1). Phylogenetic analysis revealed that the proteins encoded by the *At1g26660* and *At1g49245* genes were the likely orthologs of the UXT and ASDURF prefoldin-like proteins from animals, respectively (Fig. S2A,B). This result suggested the existence of the PFDL complex in *Arabidopsis*. To confirm this, we performed a reciprocal AP-MS using extracts from PSB-D cell suspensions expressing GS-UXT (Fig. S1). We identified AtURI, PFD2, PFD6, PDRG1, and ASDURF as top interactors of GS-UXT (Table 1; Table S1). With all likelihood, these results confirm the presence of the PFDL complex in *Arabidopsis* and that AtURI is one of its subunits.

**Table 1.** Interactors of AtURI and UXT identified by AP-MS. None of these proteins were identified in the control GS line. *Peptides*, unique peptides; *R*, biological replicates.

|  | GS-AtURI |  |  |  |  |  | GS-UXT |  |  |  |  |  |
| --- | --- | --- | --- | --- | --- | --- | --- | --- | --- | --- | --- | --- |
|  | *Peptides* |  |  | *Score Sequest HT* |  |  | *Peptides* |  |  | *Score Sequest HT* |  |  |
|  | *R1* | *R2* | *R3* | *R1* | *R2* | *R3* | *R1* | *R2* | *R3* | *R1* | *R2* | *R3* |
| **AtURI** | 63 | 67 | 65 | 2050 | 1791 | 1822 | 40 | 33 | 43 | 784 | 361 | 845 |
| **UXT** | 25 | 21 | 24 | 140 | 162 | 137 | 30 | 20 | 31 | 2034 | 933 | 2222 |
| **PDRG1** | 11 | 11 | 12 | 73 | 71 | 68 | 12 | 5 | 9 | 68 | 25 | 80 |
| **ASDURF** | 11 | 12 | 10 | 55 | 54 | 36 | 5 | 4 | 8 | 22 | 8 | 28 |
| **PFD2** | 20 | 24 | 21 | 159 | 176 | 158 | 25 | 19 | 30 | 830 | 346 | 1052 |
| **PFD6** | 27 | 28 | 26 | 141 | 143 | 123 | 22 | 18 | 28 | 133 | 70 | 165 |

Interestingly, in our AP-MS experiments, we did not pull down any canonical prefoldin other than PFD2 and PFD6. This result suggests that both AtURI and UXT are part of the PFDL complex and do not appear to form alternative complexes with other canonical prefoldins. This situation is very similar to that in animals, where URI and UXT co-immunoprecipitated only PFDN2 and PFDN6 among the canonical prefoldins [34]. However, in animals, ASDURF and PDRG1 have been suggested to be part of alternative PFD complexes, with ASDURF and PDRG1 replacing PFDN1 and PFDN4, respectively [5,34].

### Predicted structure of the Prefoldin-like complex

Although the structure of the PFDL complex is not known for any organism, the structure of the complex in humans has been predicted based on the structure of the canonical PFD complex [5]. Therefore, we wanted to determine whether the subunits of the PFDL complex from *Arabidopsis* could arrange themselves in a similar structure. The canonical PFD complex consists of two types of subunits, α-type, in which two β-hairpins connect two α-helices, and β-type, in which one β-hairpin connects the two α-helices [35]. Prediction based on modeling showed that the *Arabidopsis* PFD2 and PFD6 can adopt the structure of β-type prefoldins [24], and the AtURI prefoldin domain that of an α-type [6]. We modeled the structures of *Arabidopsis* UXT, PDRG1, and ASDURF based on the structure of predicted *Arabidopsis* PFD5, PFD4, and PFD1 [24], respectively, assuming that there is a parallelism in the arrangement of subunits in the canonical and PFDL complexes, as in humans [5]. *Arabidopsis* UXT adopted an α-type structure, whereas PDRG1 and ASDURF adopted a β-type structure (Fig. S3A-C). Using the predicted structures and the arrangement of the predicted PFDL complex from humans (Fig. S3D) [5], we could assemble the PFDL complex from *Arabidopsis* (Fig. 1A). The structural similarity between *Arabidopsis* and human prefoldins and prefoldin-like proteins strongly suggests that the *Arabidopsis* PFDL complex adopts a jellyfish-like structure *in vivo* that resembles the canonical one.

**Fig.1.**
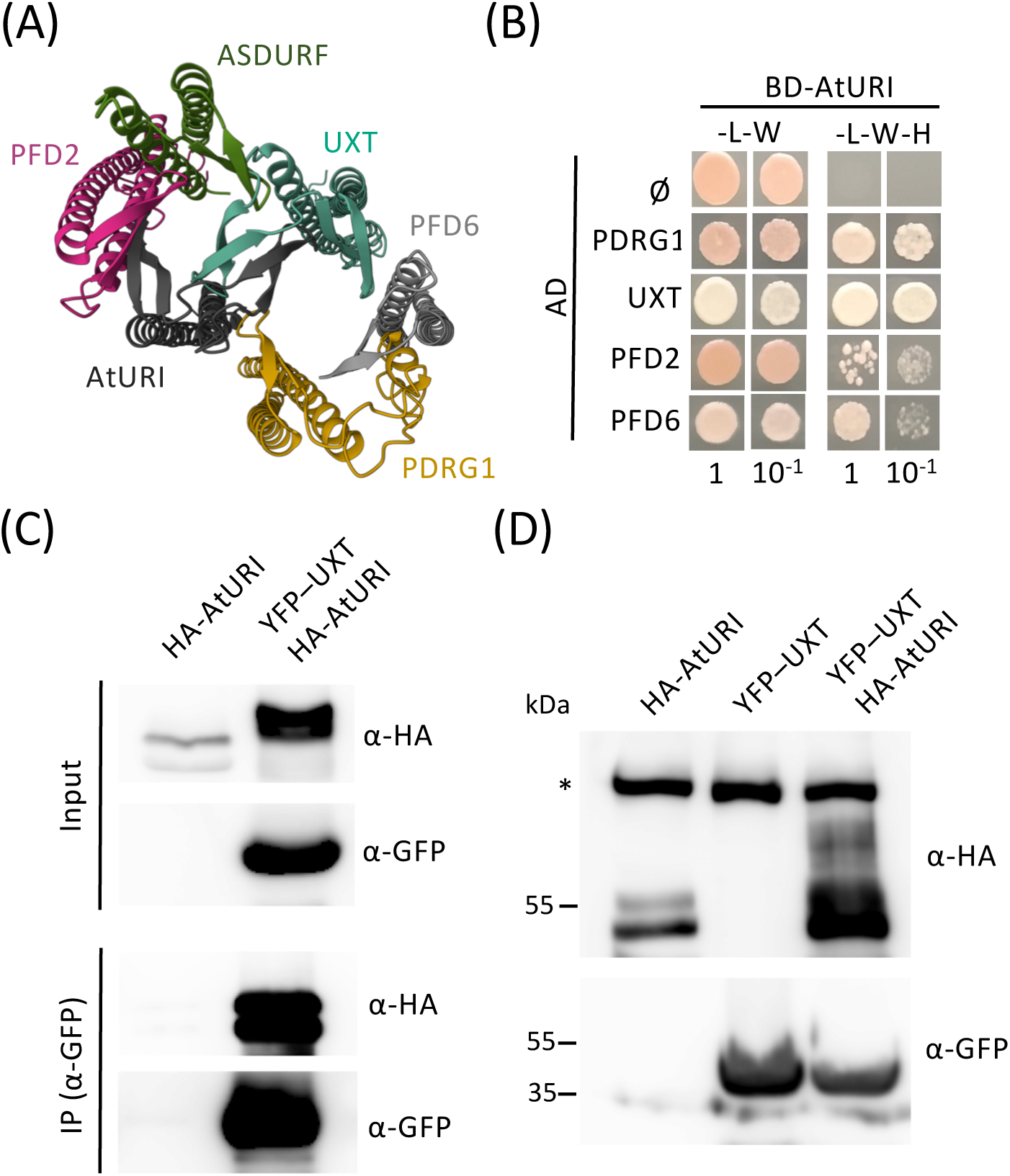
AtURI is part of the *Arabidopsis* PFDL complex. (A) Predicted 3D structure of the *Arabidopsis* PFDL complex. (B) Drop assay showing the interactions of AtURI with other subunits of the complex by Y2H. L, leucine; W, tryptophan; H, histidine. Numbers indicate the dilution used in the assay. (C) Co-immunoprecipitation assay demonstrating the interaction between HA-AtURI and YFP-UXT transiently expressed in leaves of *N. benthamiana*. Immunoprecipitations were carried out with anti-GFP coated paramagnetic beads. (D) Western analysis of protein levels of HA-AtURI and YFP-UXT transiently expressed in leaves of *N. benthamiana*. Asterisk in the upper blot marks a non-specific band that serves as loading control.

### Interactions of AtURI within the Prefoldin-like complex

In the predicted PFDL complex, AtURI occupies a central position together with UXT as α-type subunits, surrounded by the four β-type subunits. We next examined by Y2H the direct interactions of AtURI with other subunits. In agreement with the arragement in the predicted structure of the PFDL complex, AtURI interacted with the other α-type subunit, UXT, and with the two closest β-type subunits, PFD2 and PDRG1 (Fig. 1B) [6]. As reported, AtURI also interacted with PFD6 (Fig. 1B) [6], although the two subunits do not appear to occupy adjacent positions within the complex (Fig. 1A). This result could indicate that the interactions between the β-hairpins of the subunits within the complex are more extensive than assumed in the predicted model. However, it may also suggest that AtURI and PFD6 have the ability to interact with each other outside the PFDL complex.

We next examined the direct interaction between the two α-type subunits, AtURI and UXT, *in vivo*. To this end, we transiently expressed HA-AtURI and YFP-UXT in leaves of *Nicotiana benthamiana* and assessed the interaction by co-immunoprecipitation (Fig. 1C). We were able to efficiently immunoprecipitate HA-AtURI with anti-GFP antibodies from leaf extracts co-expressing both proteins, indicating that they are able to establish strong interaction *in vivo*. Interestingly, we found a higher level of HA-AtURI when co-expressed with YFP-UXT (Fig. 1C,D). This effect, however, was not observed in YFP-UXT, whose accumulation was not affected by coexpressing HA-AtURI (Fig. 1D). These latter results pointed to AtURI being an unstable protein that stabilizes upon interaction with a partner (see below).

### AtURI has an intrinsically disordered region

The interactions of URI with other subunits of the complex occur through its PFD domain in humans [36]. In *Arabidopsis*, the G-to-R change at position 90, located in the PFD domain, caused by the *aturi-1* mutation, reduces the strength of the interaction with PFD6 [6], suggesting that the PFD domain also plays a role in the interaction in *Arabidopsis*. In contrast with the other prefoldins or prefoldin-like, URI is a larger protein with a C-terminal extension after the PFD domain. We obtained the predicted structure of AtURI using AlphaFold (Fig. S4A) [37]. The algorithm predicted the position of residues that form the PFD domain with high confidence (dark blue regions in Fig. S4A and dark green areas in the Predicted Aligned Error plot in Fig. S4B), whereas the confidence relative to the position of most residues in the C-terminal extension was very low (yellow and orange regions in Fig. S4B and light green areas in the plot in Fig. S4B). Analysis of the AtURI sequence using protein disorder prediction algorithms such as FoldIndex [29], IupredL, IupredS, and SPOT-Disorder summarized in DEPICTER [27], and AlphaFold-disorder included in MobiDB [28], showed that the C-terminal extension of AtURI was predicted to be disordered (Fig. 2A,B; Fig. S4C). Therefore, AtURI can be considered a partially disordered protein, with a well-structured region at the N-terminus, the PFD domain, followed by an intrinsically disordered region (IDR).

**Fig. 2.**
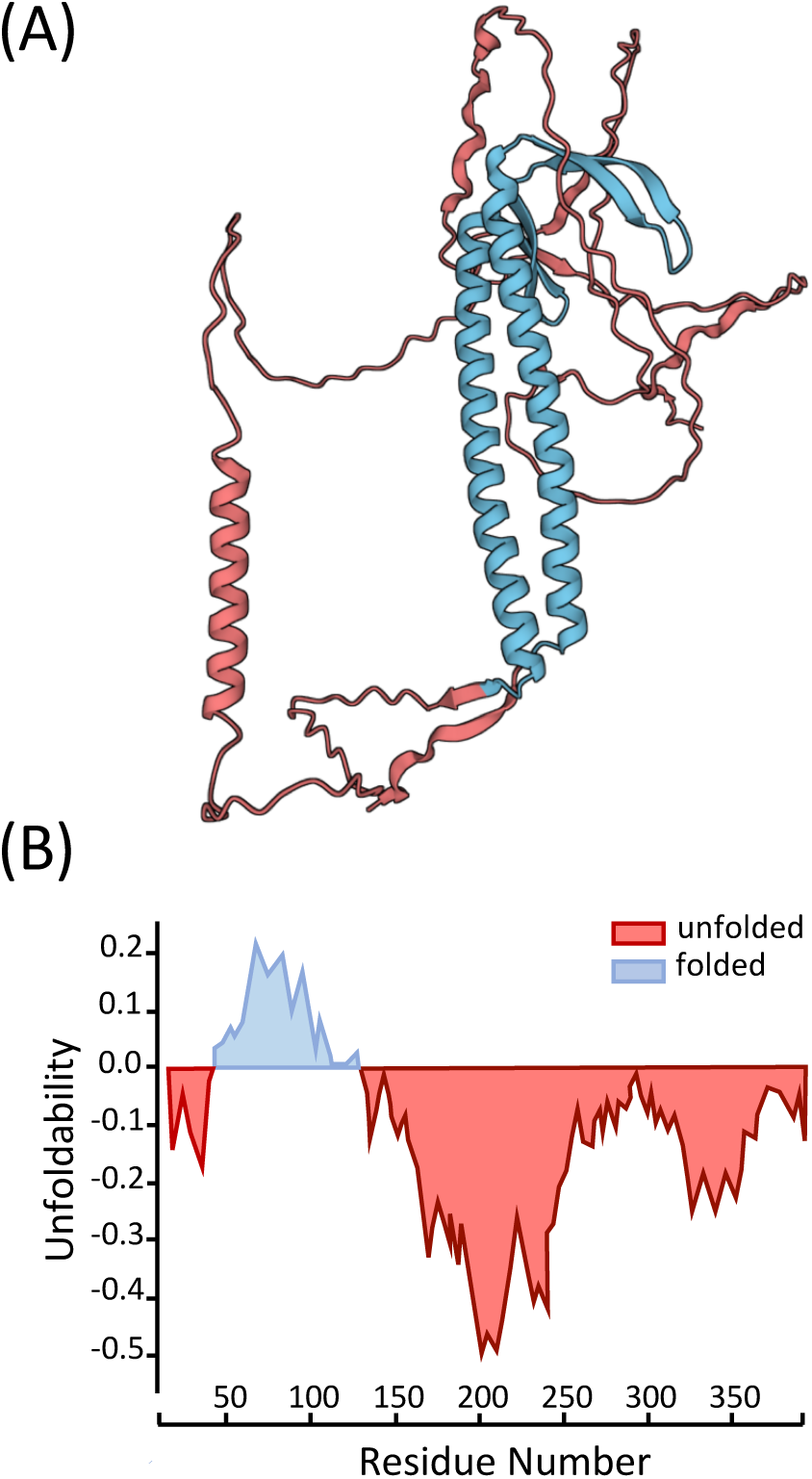
The AtURI protein is partially disordered. (A) 3D structure of AtURI predicted by AlphaFold. The part of the protein predicted to be folded is colored in blue, while the disordered/unfolded regions appear in red. (B) Output graph from the preFold analysis using the FoldIndex algorithm, showing the predicted folded (blue) and unfolded (red) regions of AtURI.

Next, we examined whether the presence of the IDR is a conserved feature of URI. We used AlphaFold and the same disorder predictors as for AtURI to analyze the human and yeast URI orthologs. The structural prediction using AlphaFold showed that human URI and yeast URI/Bud27 have a well-defined PFD domain at the N-terminus followed by a mostly unstructured region (Fig. S5A,B; Fig. S6A,B). Indeed, the disorder predictors revealed that both proteins contain an IDR spanning the region at the C-terminus of the PFD domain (Fig. S5C; Fig. S6C). Despite the low sequence homology of the C-terminal extension of URI among *Arabidopsis*, humans, and yeast [2], it is an IDR in all three cases. This suggests that the unfoldability of this part of the protein is relevant for function, while the sequence may determine species-specific features, e.g. interaction with partners.

Experimental evidence from yeast and humans has demonstrated that the region encompassing the IDR plays an important role in URI function [10,13,14,38]. For example, complementation analyses in yeast with deleted versions of URI/Bud27 have shown that the region downstream of the PFD domain, corresponding to the IDR, is sufficient to suppress the translational defects [38] and temperature sensitivity [10] of Δ*uri/bud27* cells. Next, we investigated whether AtURI exhibits characteristics of proteins with IDRs. We focused on the ability of AtURI to interact with other proteins, the stability of the AtURI protein, and the consequences for the plant of AtURI overaccumulation.

### AtURI has an extensive interactome

IDRs provide proteins with promiscuity and plasticity to interact with many different partners [39]. The ability of IDRs to provide binding promiscuity and plasticity lies in small regions called molecular recognition features (MoRFs; also called short linear motifs or linear interacting peptides) that can fold into different conformations upon binding to different interacting partners [40]. Analysis of AtURI using the fMoRFpred algorithm, included in DEPICTER [27], predicted the presence of MoRFs in the IDR, particularly in the region extending from residues 150 to 250 and at the C-terminal end (Fig. S4C). The same analysis found numerous MoRFs in the IDRs of human and yeast URI (Fig. S5C; Fig. S6C). This analysis suggests that the IDR of URI may confer it promiscuity in binding partners in the three species.

The interactome obtained by AP-MS showed that AtURI can interact *in vivo* with a relatively large number of partners, namely 135, considering the strict cutoff established for the identification of interactors, i.e. only proteins present in the three replicates, with minimum of two unique peptides in at least one of them and absent in the control line were considered (Fig. 3A; Table S1). Next, we examined whether URI is also promiscuous in yeast and humans. We used the list of URI interactors in human HEK 293T cells identified by AP-MS [5]. The list of yeast URI/Bud27 interactors, identified by various experimental approaches, was obtained from the *Saccharomyces* Genome Database [41]. URI had 142 and 117 partners in humans and yeast, respectively (Table S1) [5,41], suggesting that both are promiscuous proteins. The number of interactors, however, was smaller for the AtURI partner UXT (Fig. 3A; Table S1). In contrast to AtURI, UXT is a largely ordered protein consisting almost exclusively of the α-type PFD domain (Fig. S3A). These results suggest that promiscuity for interactions in AtURI is mediated by the IDR region rather than the PFD domain, a property likely influenced by the presence of numerous MoRFs within the IDR.

**Fig. 3.**
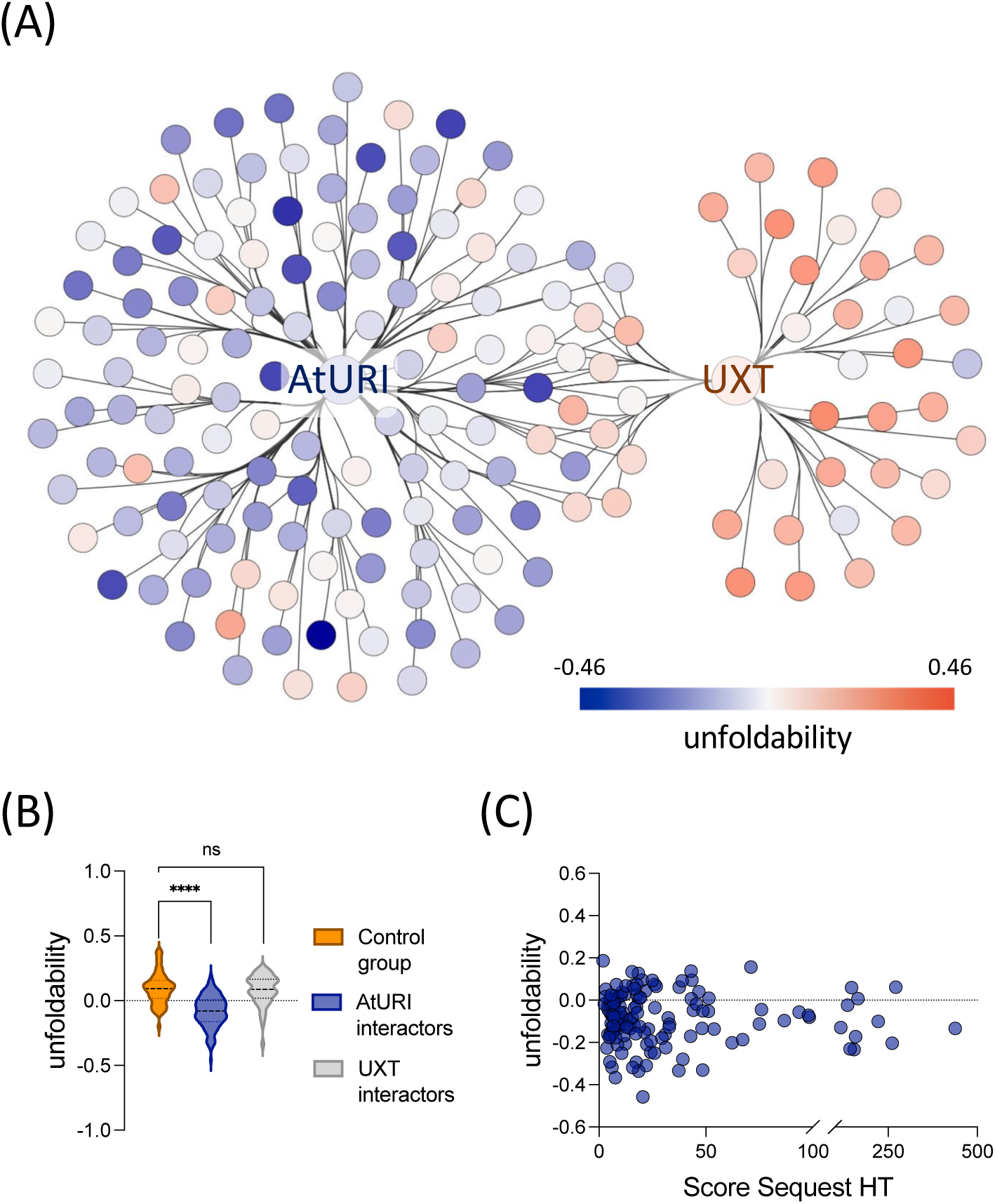
AtURI is a promiscuous protein that preferentially interacts with disordered proteins. (A) Visualization of the AtURI and UXT interacting network generated by Cytoscape. The node color indicates the unfoldability, as depicted in the scale bar. Six UXT interactors have not been included in the network, as we did not retrieve their unfoldability values from the analysis with preFold. (B) Violin plots showing the unfoldability value for a group of randomly selected *Arabidopsis* proteins (control group; n=135), AtURI interactors (n=135), and UXT interactors (n=51). Dotted lines within the violins represent the median and the first and third quartiles. ****, *P* < 0.0001; ns, non-significant. (C) Scatter plot representing the unfoldability value and Score Sequest HT for each AtURI interactor.

A higher preference for interactions between disordered proteins was found in humans [42]. To investigate whether AtURI preferentially interacts with disordered proteins, we determined the unfoldability value for each of the AtURI interactors using FoldIndex and compared it with the value calculated for a list containing the same number of randomly selected proteins (Table S2). Positive values correspond to preferentially folded proteins, while negative values correspond to predicted disordered proteins; the lower the value, the more disordered [29]. For example, the unfoldability value of SERRATE (SE), one of the AtURI interactors with features of disordered proteins [43], is -0.133, whereas the value of AtURI is -0.099. The average unfoldability value of the AtURI interactors was negative and significantly lower than that of the control group, which was positive (Fig. 3A,B), indicating that there were more proteins among the AtURI interactors predicted to be disordered or to have IDRs. Next, we investigated whether this property of AtURI is conserved in humans and yeast or, on the contrary, whether it is species dependent. As for AtURI, we obtained the unfoldability value for each URI partner in each species using FoldIndex. In contrast to AtURI, the average unfoldability value for the human URI interactors was similar to the control list, in both cases positive (Fig. S7 and Table S2). Meanwhile, the average unfoldability value of URI/Bud27 interactors was positive but significantly lower than the control list (Fig. S7 and Table S2). Our analysis suggests that the composition of the URI interactomes probably depends on the species-specific sequence of URI, including the IDR, and is not solely determined by the IDR unfoldability.

Next, given the prefered interaction of AtURI with disorderd proteins, we sought to determine whether the degree of unfoldability of each partner had an impact on the strength of the interaction. To this end, we plotted the unfoldability value against the average score Sequest HT [44] for each of them in the three replicates (Table S3), which we used as a proxy to measure the strength of the interaction. As can be seen in Fig. 3C, there was no correlation between these two parameters, indicating that the degree of disorder of the partner does not seem to determine the strength of its interaction with AtURI.

In contrast to AtURI, the average unfoldability value of the UXT interactors was positive and similar to that of the control group of proteins (Fig. 3A,B). The specific prevalence of disordered proteins among AtURI partners suggests that AtURI establishes these interactions alone and not as part of the PFDL complex. This is corroborated by the limited overlap between AtURI and UXT interactors, as depicted in Fig. 3A, which weakens the case for AtURI’s partners being associated with the PFDL. However, it is important to note that we cannot completely exclude the possibility that AtURI interacts with its partners as part of the PFDL, and the fact that we could not identify a larger number of UXT interactors may be due to technical limitations of the immunoprecipitation.

Gene Ontology (GO) analysis of AtURI partners showed that the most enriched categories in “Molecular Function” were related to RNA binding, nucleic acid binding, and mRNA binding, while the most enriched categories in “Biological Process” were related to mRNA processing, mRNA metabolic process, and RNA splicing (Table S4). This observation is in agreement with studies showing that RNA-binding proteins and components of the spliceosome are largely disordered [45–47] and suggests a functional relationship between AtURI and RNA transactions based on the disorder of AtURI and its RNA-binding partners.

### AtURI is an unstable protein likely degraded via 20S proteasome

The involvement of disordered or partially disordered proteins in regulatory pathways, as well as their promiscuity in establishing protein-protein interactions, means that the amount of these proteins must be tightly controlled [19]. Thus, disordered or partially disordered proteins tend to be unstable, short-lived proteins [19]. We investigated whether this was the case for AtURI by conducting *in vitro* cell-free degradation assays. To this end, we transiently expressed YFP-AtURI and GFP in leaves of *N. benthamiana* and incubated the protein extracts at 30 °C for several time points in the presence of the protein synthesis inhibitor CHX. YFP-AtURI and GFP protein levels were analyzed in Western blots. As shown in Fig. 4A-D, levels of YFP-AtURI decreased within the first minutes of incubation, while the GFP levels were not changed. This result suggests that AtURI is an unstable protein. Next, we investigated whether AtURI is degraded via the proteasome. Incubation of the protein extracts with the proteasome inhibitor MG-132 delayed the degradation of YFP-AtURI, suggesting that it is at least partially mediated via this pathway (Fig. 4A,B).

**Fig. 4.**
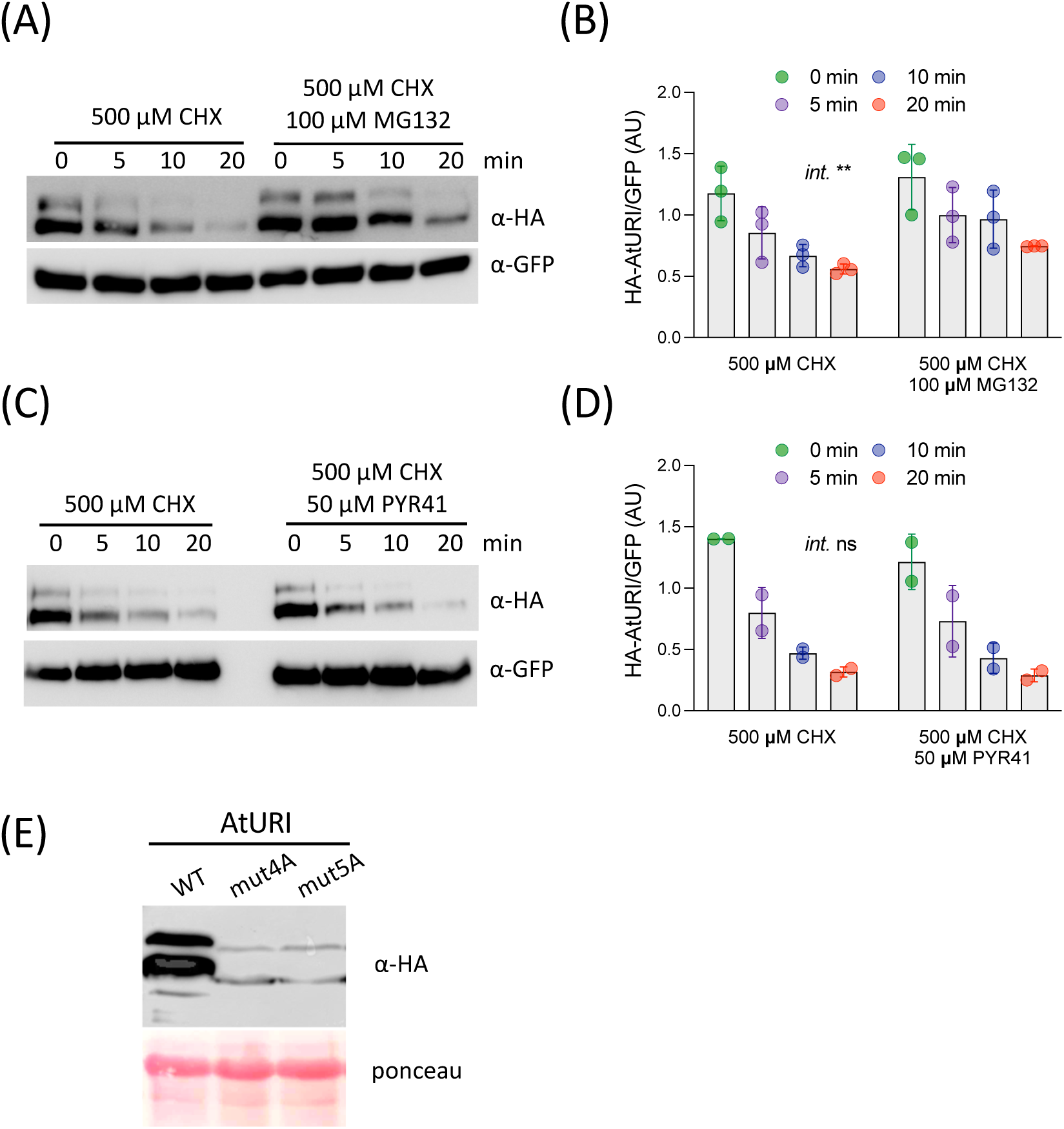
AtURI is an unstable protein. Total proteins from *N. benthamiana* leaves co-expressing HA-AtURI and GFP were used for cell-free degradation assays. Protein extracts were incubated with 500 μM CHX or with 500 μM CHX and 100 μM MG-132 (A, B), or with 500 μM CHX and 50 μM PYR-41 (C, D) for the indicated times. HA-AtURI and GFP levels were determined from three (A, B) and two (C, D) biological replicates by western blot. One representative western blot is shown in (A, C), while the quantifications are shown in (B, D). AtURI normally appears as a double band; the lower band was used for quantification. (E) Western blot showing the protein level of the WT and mutant versions of HA-AtURI transiently expressed in leaves of *N. benthamiana*. In (B, D) *Int.* refers to the interaction between treatments assessed by two-way ANOVA; **, *P* < 0.01, ns, non-significant.

Many disordered proteins are directly degraded by the 20S core proteasome without requiring the proteasome 19S lid, which is responsible for unfolding the target proteins [48]. This process is independent of protein ubiquitination. The 20S proteasome also degrades partially disordered proteins, as shown for the *Arabidopsis* protein SE [43]. To determine whether AtURI can also be degraded via 20S proteaseome, we conducted the *in vitro* cell-free degradation assays in the presence of CHX and PYR-41, an inhibitor of the ubiquitin-activating enzyme E1. As shown in Fig. 4C and D, the decay of YFP-AtURI protein levels was not affected by PYR-41, suggesting that it occurs independent of ubiquitination via 20S proteasome. The degradation of the AtURI protein was also observed when we conducted the *in vitro* cell-free degradation assays with recombinant AtURI fused to maltose binding protein (MBP) (Fig. S8). The degradation of the recombinant MBP-AtURI was delayed by treatment with MG-132 but not with PYR-41, suggesting that, like AtURI expressed in *N. benthamiana* leaves, the recombinant version was degraded via the core 20S proteasome.

IDRs are specially accessible to post-translational modifications such as phosphorylation, which in many cases alter the stability of the protein [16]. Phosphoproteomic analyses identified 13 *in vivo* phosphorylated Ser/Thr residues in AtURI [49–51], all in the IDR (Fig. S4C; Table S5). We tested whether the phosphorylation of some of these residues had an effect on the protein level of AtURI. We prepared two mutant versions of AtURI in which two clusters of phosphorylated Ser/Thr were replaced by Ala. The replaced residues were Ser^165^, Ser^174^, Thr^178^, Ser^211^, and Ser^218^ in AtURI^mut5A^, all of them predicted to be phosphorylated by Casein Kinase II (CK2; Table S5), and Ser^262^, Ser^263^, Ser^264^, and Ser^268^ in AtURI^mut4A^. We transiently expressed the WT and mutant versions fused to YFP in leaves of *N. benthamiana* and analyzed the protein levels by Western blot (Fig. 4E). Two interesting findings could be derived from this analysis. First, the bands corresponding to the mutant versions migrated faster than those of the WT version, suggesting that the mutated residues were normally phosphorylated in the leaves of *N. benthamiana*. Second, the mutant proteins accumulated to a lesser extent than the WT counterpart, suggesting that phosphorylation of these residues may play a role in maintaining the stability of AtURI. It is tempting to speculate that CK2, a Ser/Thr kinase conserved in eukaryotes with high and constitutive activity [52], is one of the kinases that help maintain AtURI phosphorylated and stable, as several *in vivo* phosphorylated Ser/Thr on AtURI have been reliably predicted as target sites of CK2 (Table S5). It has been proposed that the interaction with partners can stabilize disordered proteins [43]. The phosphorylation of residues within the IDR could enhance AtURI’s interaction with its partners, thereby contributing to its stabilization. This could occur by a mechanism involving phosphorylation-induced folding of the disordered region that mediates the interaction [53]. Alternatively, it may directly influence stability by preventing AtURI from being targeted for degradation by the proteasome. An example related to the latter scenario, albeit resulting in the opposite outcome, is the phosphorylation of the partially disordered protein SE by PRP4KA in *Arabidopsis*, which triggers the degradation of SE [54].

### Overaccumulation of AtURI interferes with its function

To investigate the effects of AtURI overaccumulation on the plant, we prepared several transgenic *Arabidopsis* lines expressing a *YFP-AtURI* fusion under the control of the strong promoter *35S*. These lines exhibited varying degrees of developmental and growth changes, some of which were observed in more than one line (Fig. 5). These included the appearance of twin stems (Fig. 5A), slight changes in phylotaxis (Fig. 5B), and severe dwarfism and bushy appearance, probably due to internode shortening with additional branching (Fig. 5C,D). The phenotype of homozygous plants of lines 4.9, 6.7, 16.11 and 29.6 was very similar within each line. However, a range of phenotypes was observed in homozygous plants of line 5.3 (Fig. 5A,C,D). To determine whether the severity of the phenotypes was directly related to the amount of YFP-AtURI, we measured it in inflorescences of 25-day-old plants (Fig. 6A). Surprisingly, the dwarf lines 5.3 and 6.7 had the lowest YFP-AtURI levels. We ruled out that these phenotypes were caused by co-supression of the endogenous *AtURI*, as it was overexpressed in these lines and the transgene was expressed (Fig. S9A,B). Reduced level of YFP-AtURI was also observed in 7-day-old seedlings of the line 5.3 (Fig. 6B), with no correlation between the amount of YFP-AtURI protein and transgene expression (Fig. 6B; Fig. S9C). None of the transgenic lines showed a distinctive phenotype at the seedling stage (Fig. S9D).

**Fig. 5.**
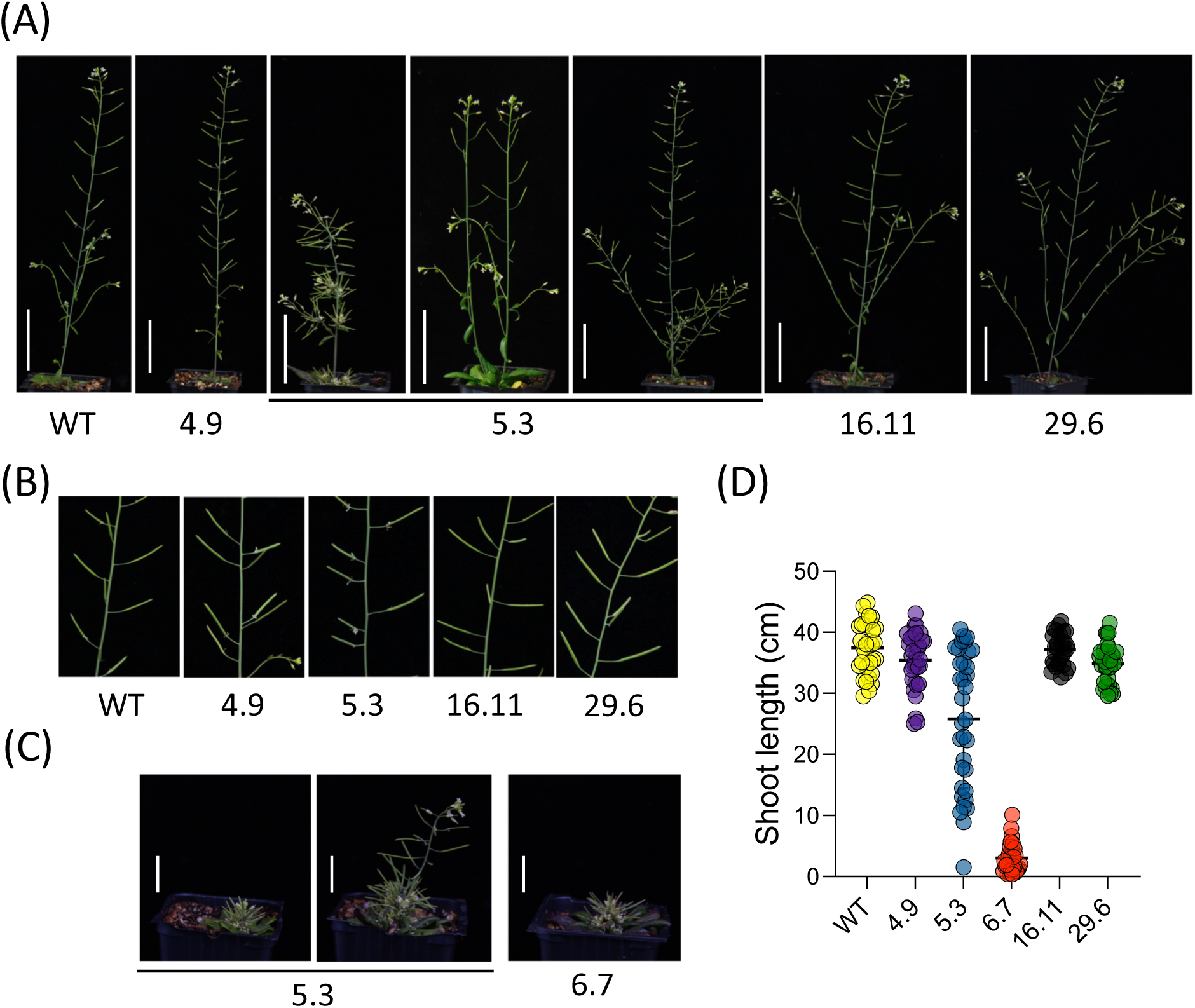
Alteration of AtURI levels causes growth and developmental defects. Representative homozygous plants from the indicated lines expressing *35S:YFP-AtURI* are shown in (A-C). Plants were grown under a 16 h light:8 h dark photoperiod at 22 °C for 25 days. Several plants of line 5.3 are shown, as this line exhibited phenotypes of diverse severity. (B) Close-up view of the defects in silique arrangement in the transgenic lines. (C) Dwarf and bushy plants from lines 5.3 and 6.7. (D) Stem length of 25-day-old plants from the different lines; 28-35 plants per line were used for measurements. The scale bar in (A) and (C) is 5 cm and 2 cm, respectively.

**Fig. 6.**
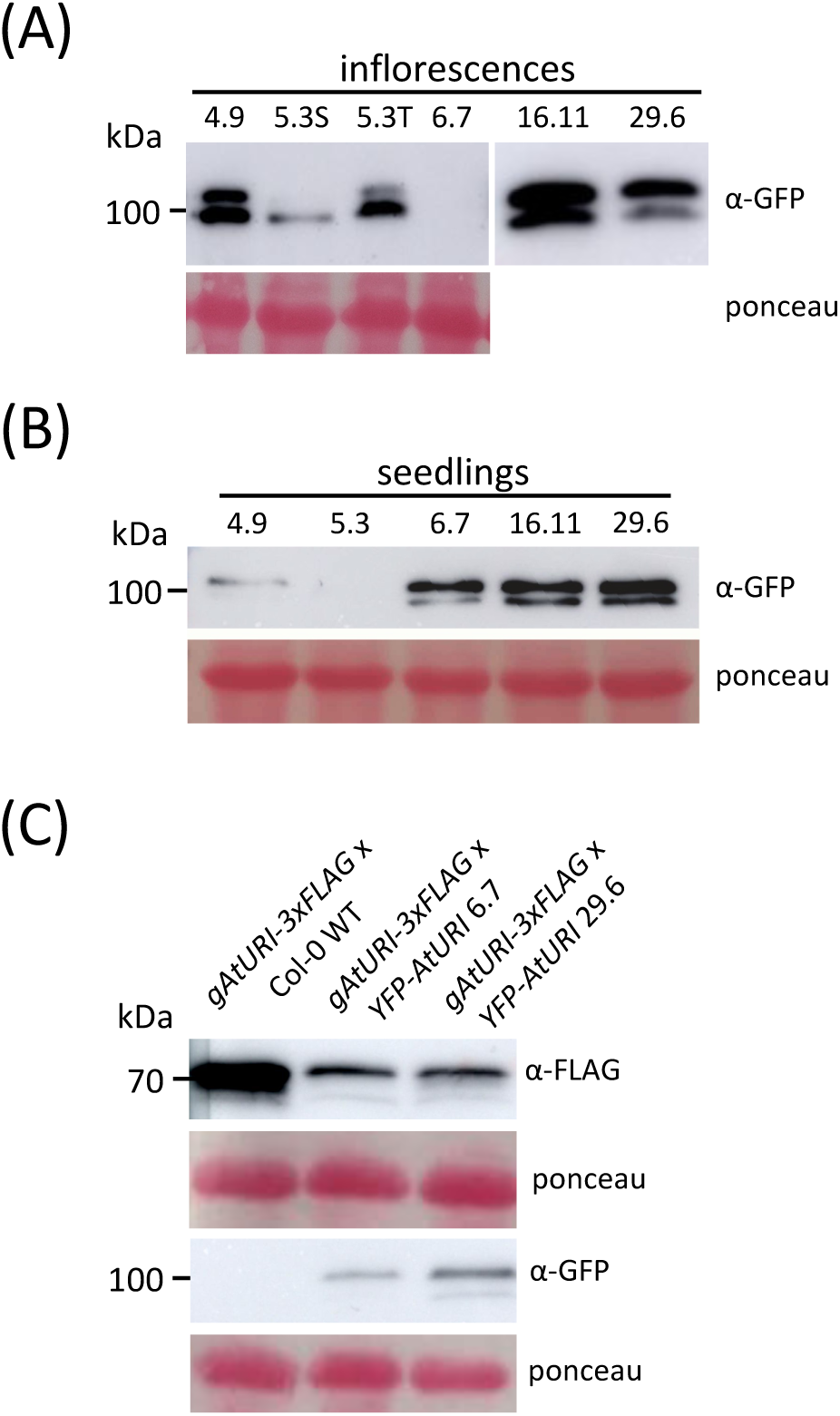
Overexpression of *YFP-AtURI* causes AtURI degradation. Western blot analysis of YFP-AtURI levels in inflorescences of 25-day-old plants (A) and 10-day-old seedlings (B) of the *35S:YFP-AtURI* transgenic lines. In (A), 5.3T and 5.3S refer to the inflorescences of tall and dwarf plants, respectively. (C) Western blot analysis of AtURI-3xFLAG and YFP-AtURI levels in 10-day-old seedlings of the indicated genotypes. Ponceau staining is shown as a loading control.

A similar situation was reported for SE [43]. The excess of transgenic SE protein in the *SE* overexpressing lines triggered its degradation by the 20S proteasome, which also degraded the endogenous SE resulting in a similar phenotype of *se* mutants and *SE* transgenic lines. Li et al. (2020) proposed that SE is normally part of different protein complexes and any fraction of SE not associated with a complex is degraded by the 20S proteasome to prevent undesirable effects of the excess of this partially disorderd protein. SE, like other proteins with disordered regions, can be protected from degradation by masking the unfolded region if it is part of a complex, or by undergoing a folding-upon-interaction transition [55]. We considered that a similar mechanism could operate to control AtURI protein levels. We explored this possibility by investigating whether overexpression of *YFP-AtURI* could trigger the degradation of AtURI-3xFLAG in *pAtURI:AtURI-3xFLAG* x *35S:YFP-AtURI* 6.7 and 29.6 F1 seedlings. As shown in Fig. 6C the levels of AtURI-3xFLAG were reduced in F1 seedlings co-expressing YFP-AtURI compared to control F1s from the cross with the non-transgenic WT. This would be consistent with the reduced levels of AtURI-3xFLAG being caused by the degradation triggered by the overexpression of YFP-AtURI, which is also likely to affect endogenous AtURI. Thus, AtURI levels may be subject to a control mechanism similar to that of SE [43]. This suggests that some of the developmental phenotypes shown in Fig. 5, such as dwarfism and extra branching, were caused by reduced AtURI. These developmental phenotypes, however, are not observed in the *aturi-1* mutant [6]. This likely reflects the different molecular nature of the defects, i.e. a change in one amino acid in the aturi-1 protein compared to a reduced level of the protein in the transgenic plants, which have a different effect on the developmental processes involving AtURI.

## CONCLUSION

Our results show that the PFDL complex is present in *Arabidopsis* and that AtURI is one of its subunits. AtURI exhibits features of intrinsically disordered proteins thanks to the IDR located downstream of the PFD domain. These features include promiscuity to establish protein-protein interactions and instability of the protein. AtURI instability could allow the plant to precisely control the amount of AtURI so that isolated protein that is not incorporated into functional protein complexes is degraded to prevent unwanted effects. These properties could be reflected in the activity of AtURI, both as part of the PFDL complex and as an independent subunit. Our interactome analysis suggests a possible role for AtURI in RNA metabolism, likely exerted as an independent subunit via the promiscuity of the IDR. AtURI could also recruit interactors to the complex via the IDR and limit the formation of the complex under those physiological conditions where AtURI becomes unstable and thus limiting. We hypothesize that the instability of AtURI is regulated by environmental changes and endogenous signals via phosphorylation and dephosphorylation of residues in the IDR, which likely influences the interaction of AtURI with its partners. This regulation of AtURI could potentially serve as a mechanism to control downstream pathways, both dependent and independent of the PFDL complex.

## ACKNOWLEDGEMENTS

We thank Drs. Miguel Blázquez (IBMCP, Valencia, Spain) and Noel Blanco-Touriñán (University of Lausanne, Switzerland) for critically reviewing the manuscript, and Dr. Javier Forment (IBMCP, Valencia, Spain) for his help with the bioinformatics analysis. Research was funded by grant PID2019-109925GB-I00 from MCIN/AEI/10.13039/501100011033 to D.A. Y.G-M., A.P-A. and L.H-V. were supported by Ministerio de Economía y Competitividad (BES-2017-081041), Ministerio de Educación (FPU17/05186) and Generalitat Valenciana (GRISOLIAP/2018/001) predoctoral contracts, respectively. C.C-R. was supported by a Juan de la Cierva-Formación post-doctoral contract (FJC2020-045099-I) from the Spanish Agencia Estatal de Investigación.

