## Supplemental Figures and Figure Legends for "The prefoldin-like protein AtURI exhibits characteristics of instrinsically disordered proteins"

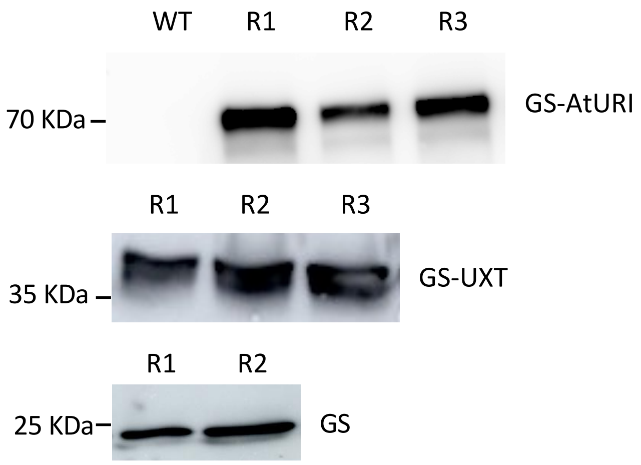


**Fig. S1.** The GS fusion proteins are expressed in transgenic *Arabidopsis* cell suspensions. Total proteins from extracts of various transgenic lines were subjected to Western analysis, and the fusion proteins were detected using a peroxidase anti-peroxidase soluble complex antibody. Protein extracts from non-transgenic cell suspensions were used as a negative control in the upper blot.

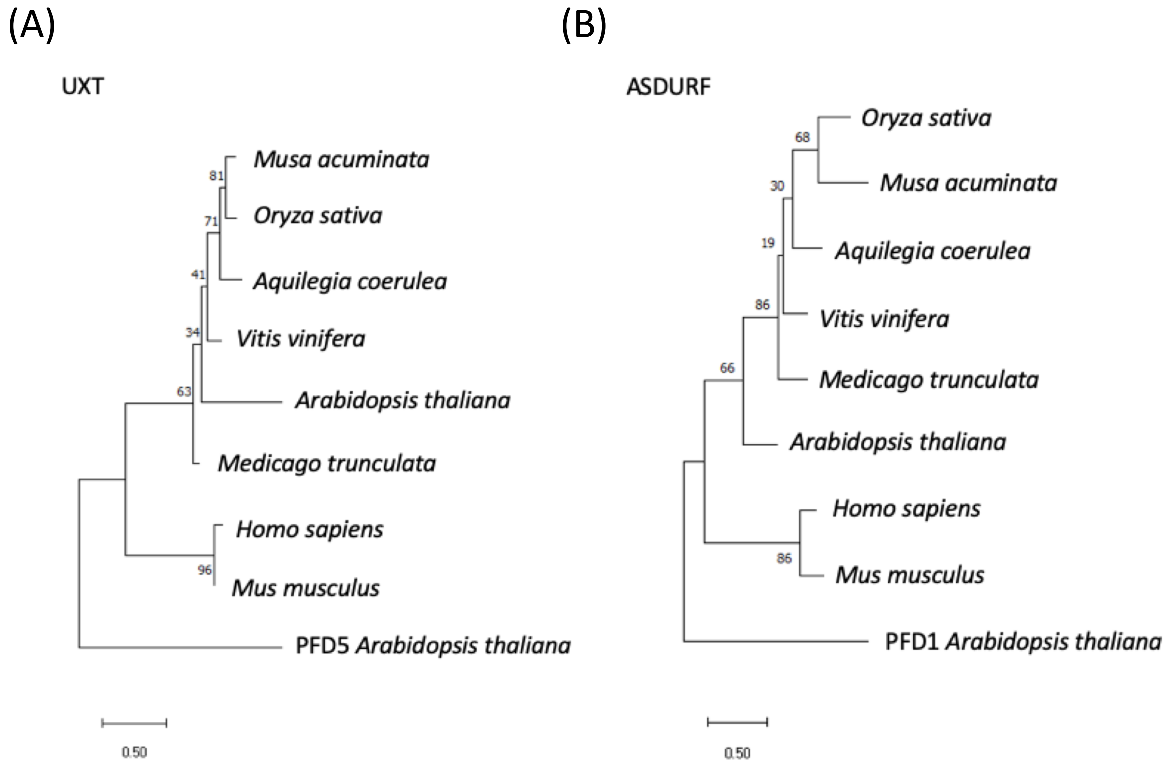


**Fig. S2.** The proteins encoded by *At1g26660* and *At1g49245* genes are the likely orthologs of UXT and ASDURF. Phylogenetic analysis of UXT (A) and ASDURF (B) proteins from various plant species, humans and mouse. Numbers in branches are maximum likelihood bootstrap values from one thousand replicates. The scale bar refers to the evolutive distance. *Arabidopsis* PFD5 and PFD1 were used as outliers.

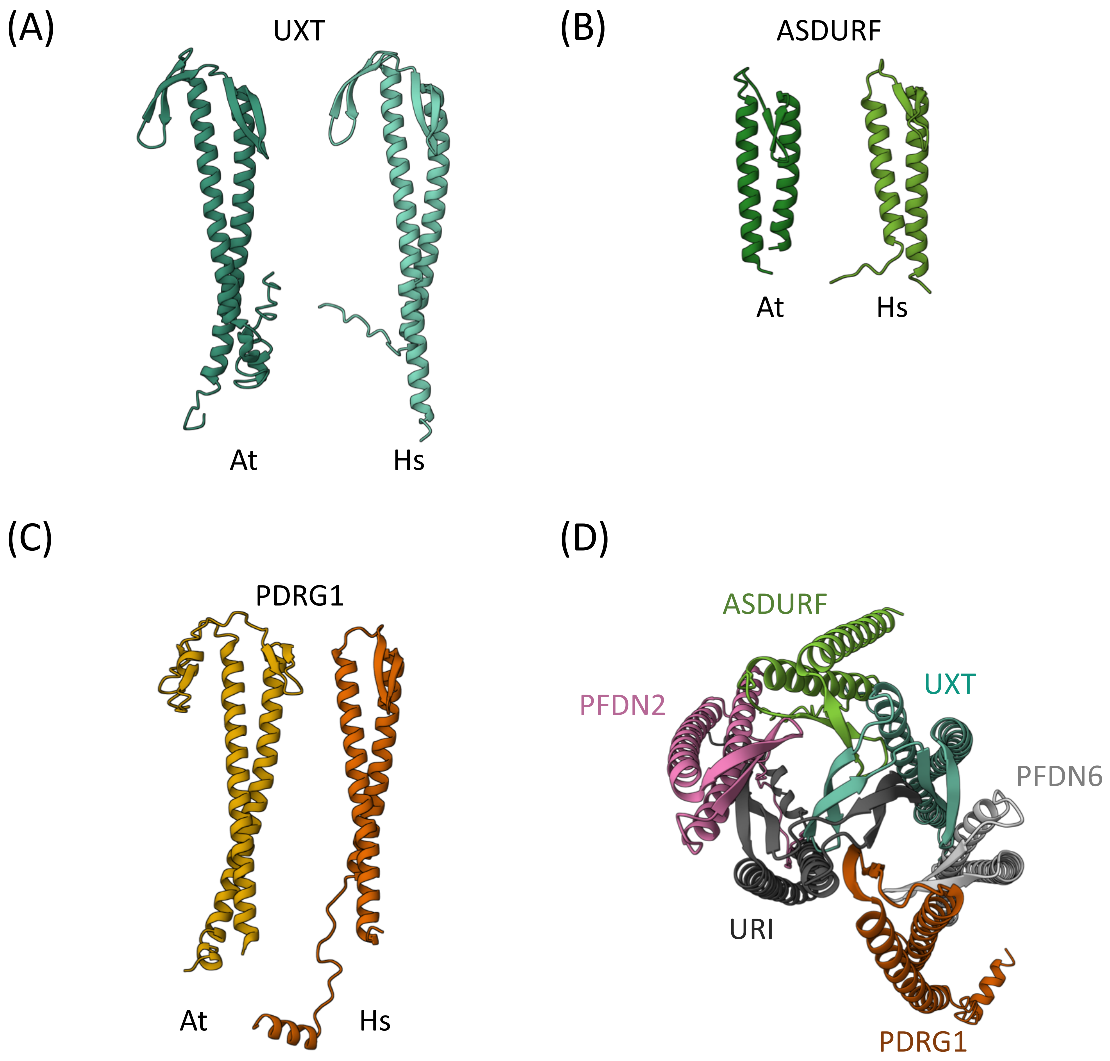


**Fig. S3.** Predicted structure of *Arabidopsis* UXT, ASDURF, and PDRG1. The structure of *Arabidopsis* (At) UXT (A), ASDURF (B) and PDRG1 (C) is shown together with the structure of their human (Hs) orthologs. (D) Predicted structure of the human PFDL complex. Only the URI PFD domain is included in the structure.

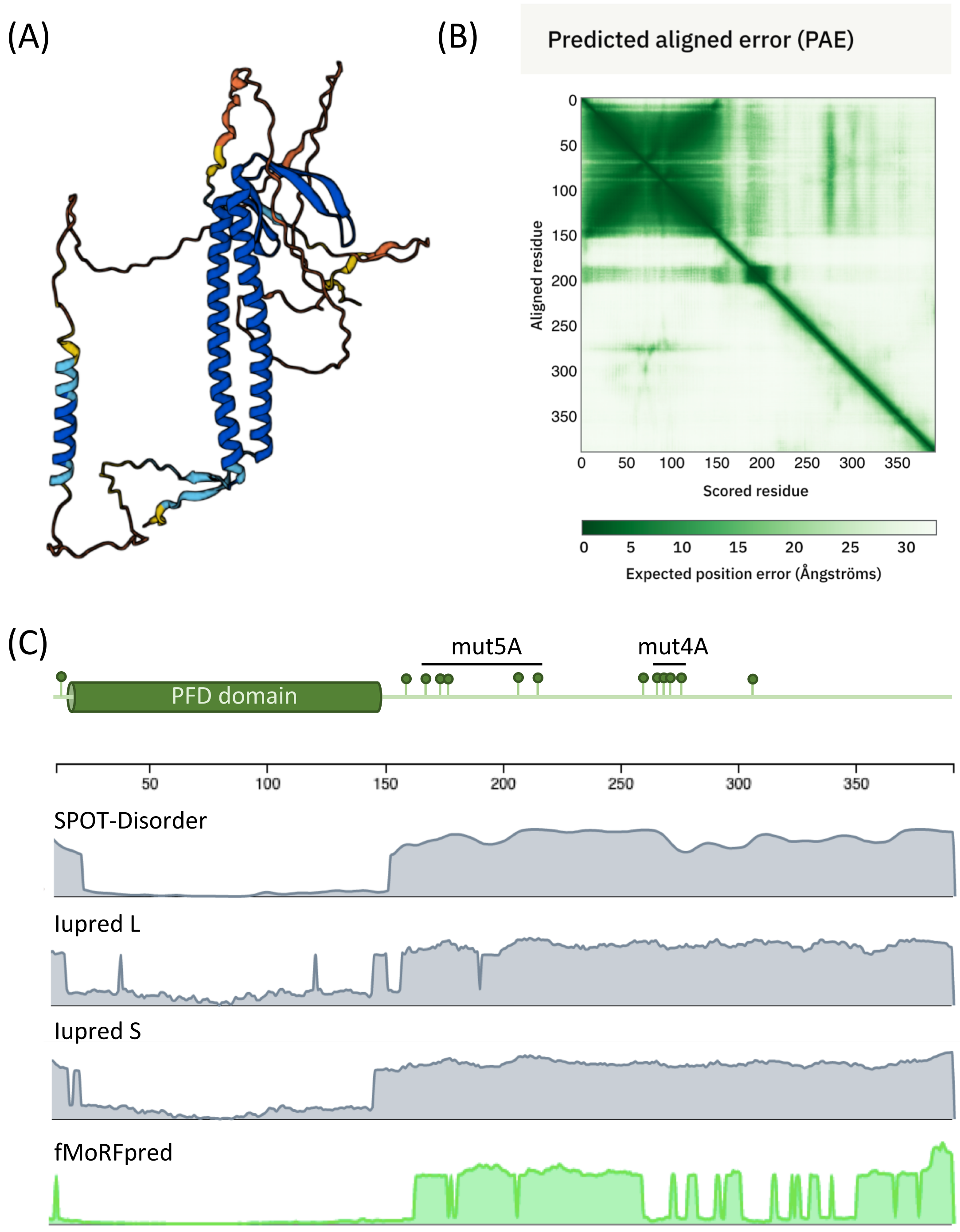


**Fig. S4.** AtURI is predicted to be a partially disordered protein. (A) The AlphaFold-predicted structure of AtURI, where colors represent model confidence: dark blue (very high), light blue (high), orange (low), and yellow (very low). (B) The PAE plot measures the confidence in the relative position of two residues in Angstroms (ranging arbitrarily between 0 and 31), with smaller distances indicating higher confidence. (C) Plots display disorder (gray) and MoRF (green) propensity profiles in AtURI predicted by SPOT-Disorder, Iupred L, Iupred S, and fMoRFpred. The numeric line at the top indicates residue positions in the protein. The scheme of AtURI is depicted at the top of the panel, with green lollypops indicating residues phosphorylated *in vivo*.

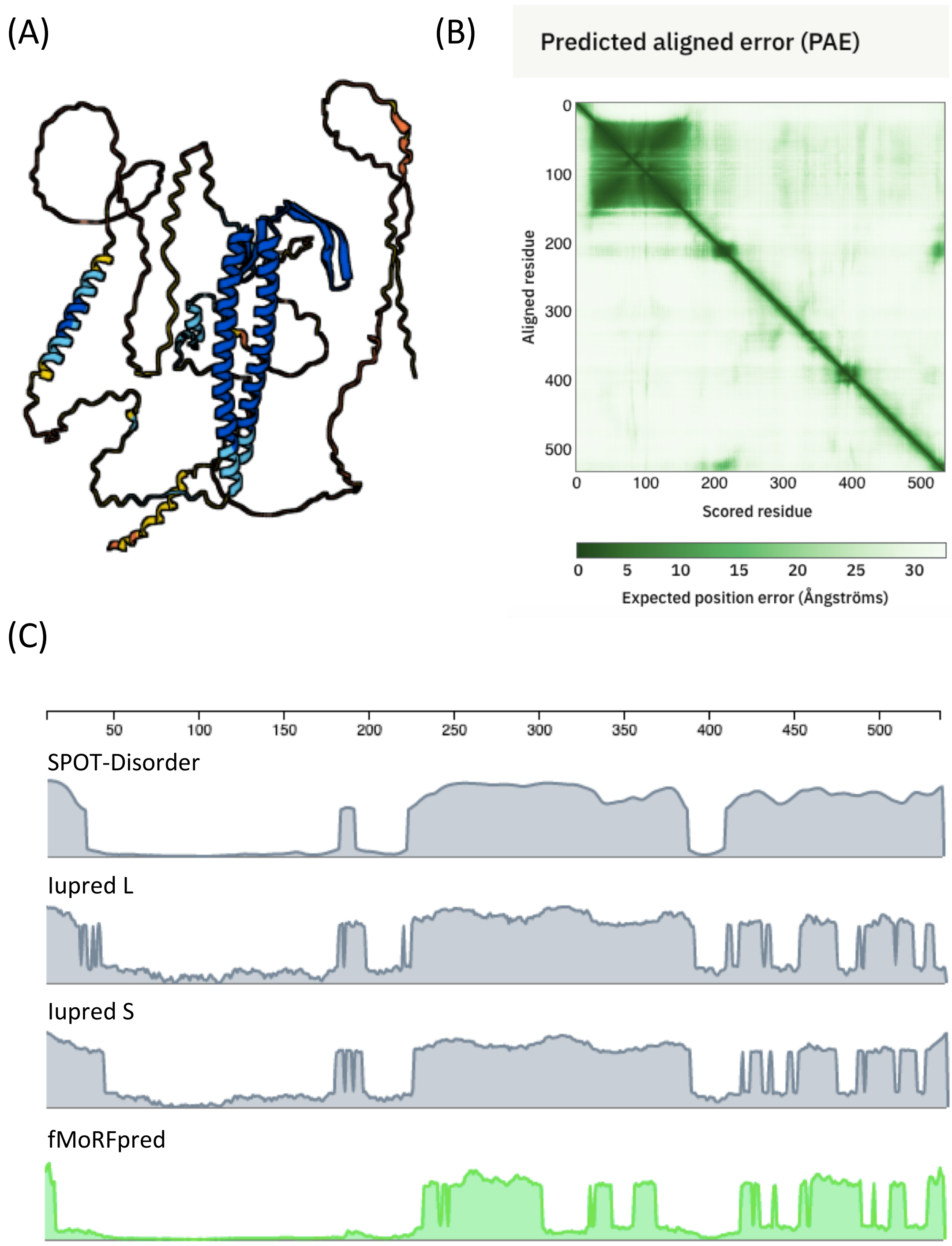


**Fig. S5.** Human URI is predicted to be a partially disordered protein. (A) The AlphaFold-predicted structure of URI, where colors represent model confidence: dark blue (very high), light blue (high), orange (low), and yellow (very low). (B) The PAE plot measures the confidence in the relative position of two residues in Angstroms (ranging arbitrarily between 0 and 31), with smaller distances indicating higher confidence. (C) Plots display disorder (gray) and MoRF (green) propensity profiles in URI predicted by SPOT-Disorder, Iupred L, Iupred S, and fMoRFpred. The numeric line at the top indicates residue positions in the protein.

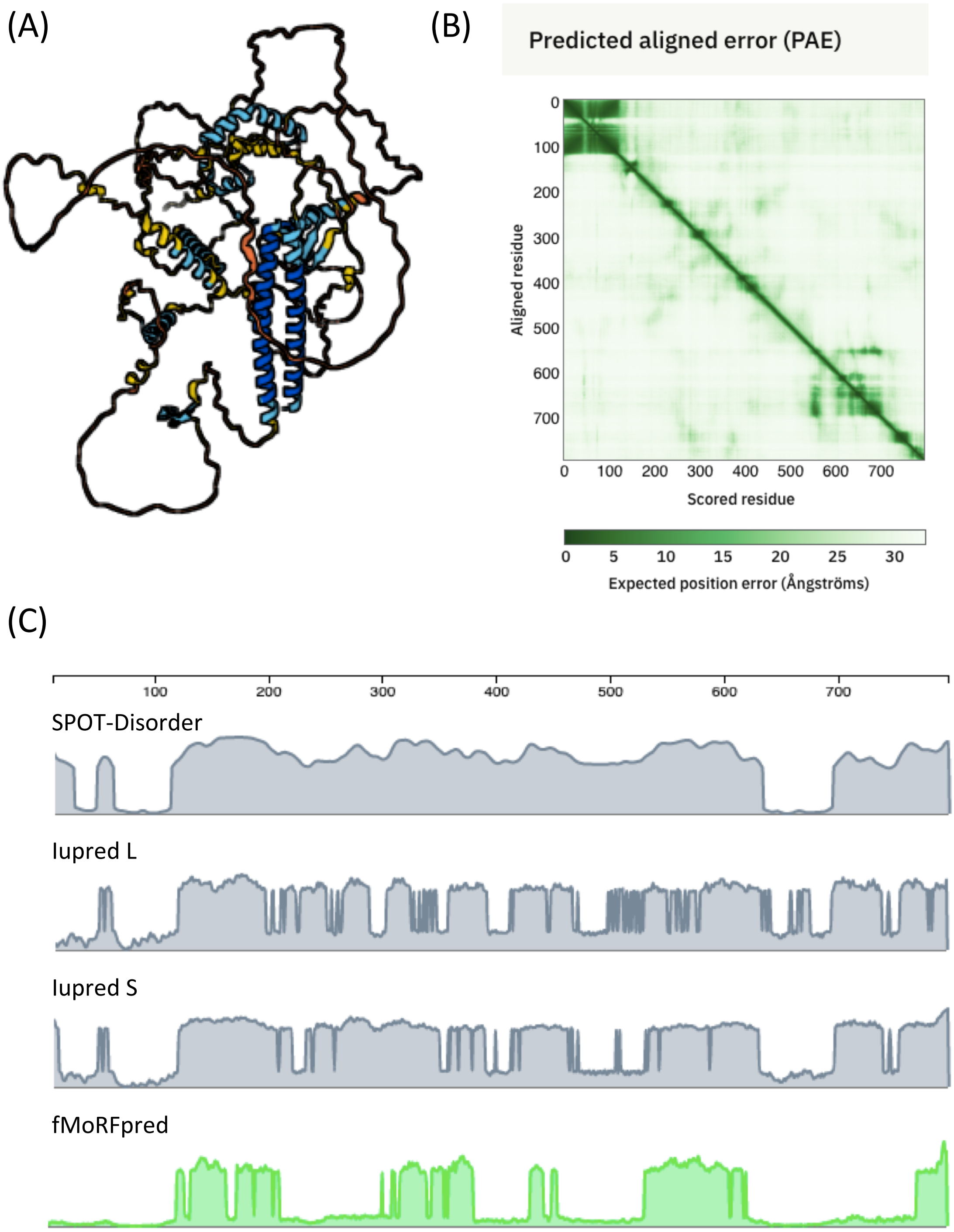


**Fig. S6.** URI/Bud27 is predicted to be a partially disordered protein. (A) The AlphaFold-predicted structure of URI/Bud27, where colors represent model confidence: dark blue (very high), light blue (high), orange (low), and yellow (very low). (B) The PAE plot measures the confidence in the relative position of two residues in Angstroms (ranging arbitrarily between 0 and 31), with smaller distances indicating higher confidence. (C) Plots display disorder (gray) and MoRF (green) propensity profiles in URI/Bud27 predicted by SPOT-Disorder, Iupred L, Iupred S, and fMoRFpred. The numeric line at the top indicates residue positions in the protein.

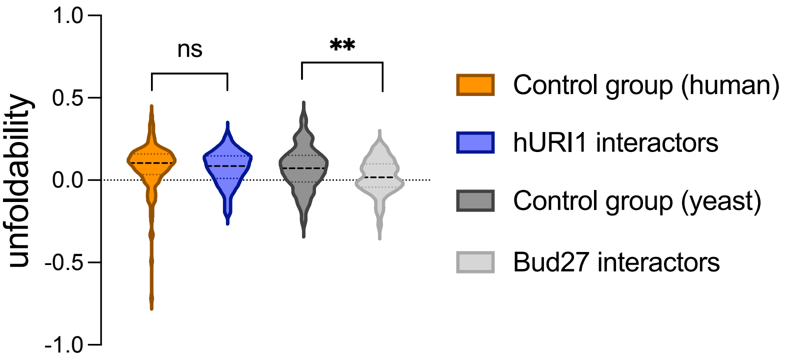


**Fig. S7.** Human URI and yeast URI/Bud27 preferentially interact with ordered proteins. Violin plots showing the unfoldability value for a group of randomly selected human (control group human; n=132) and yeast (control group yeast; n=114) proteins, human URI interactors (hURI; n=132), and yeast URI/Bud27 interactors (Bud27; n=114). Dotted lines within the violins represent the median and the first and third quartiles. **, $P < 0.01$; ns, non-significant.

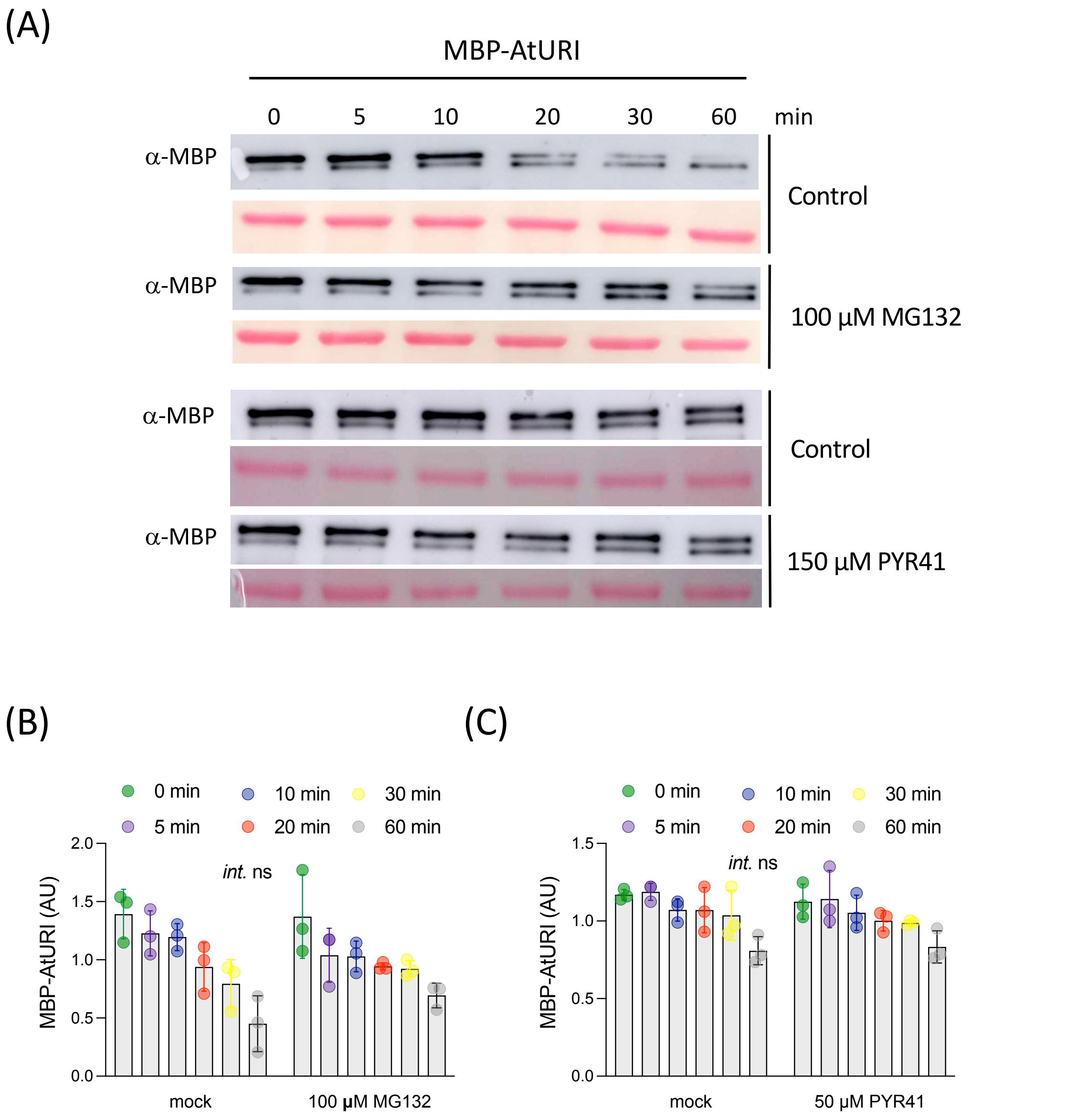


**Fig. S8.** Recombinant MBP-AtURI is likely degraded via the proteasome 20S. (A) Representative immunoblots illustrating the degradation of MBP-AtURI in cell-free degradation assays, with Ponceau staining of the membranes shown below each immunoblot. Quantification of the upper band, representing MBP-AtURI, in three replicates of mock and MG-132 treatment (B) and mock and PYR-41 treatment (C). '*Int.*' refers to the interaction between treatments assessed by two-way ANOVA. The effect of the treatments is non-significant (ns), despite a tendency towards the stabilization of MBP-AtURI observed after treatment with MG-132.

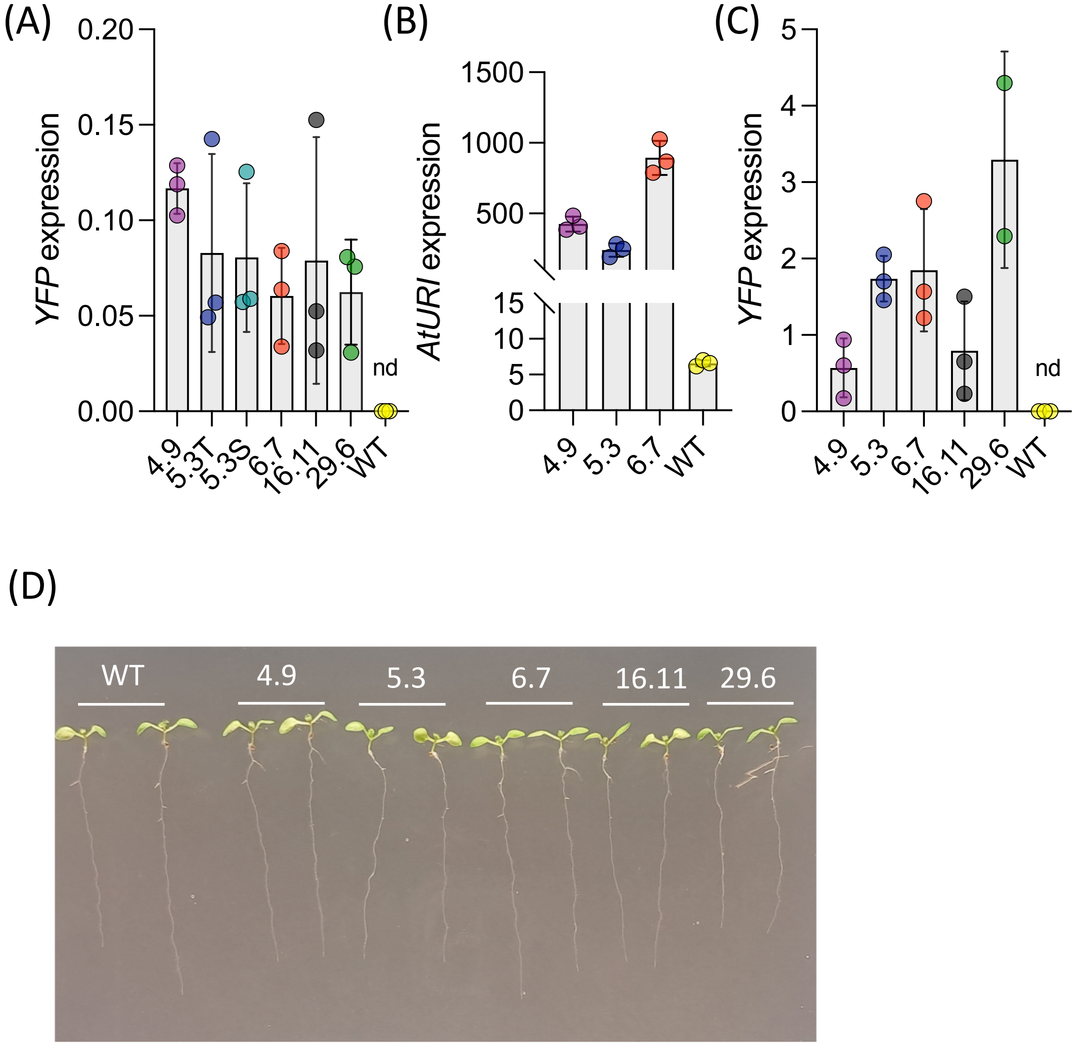


**Fig. S9.** *AtURI* is overexpressed in the transgenic lines. Expression of *YFP* (A, C) and *AtURI* (B) assessed by RT-qPCR in inflorescences (A) and seedlings (B, C). Each dot represents a biological replicate; nd, non-detected. (D) Picture of two representative seedlings of the different transgenic lines alongside the wild type.
